# Schizophrenia-derived hiPSC brain microvascular endothelial cells show impairments in angiogenesis and blood-brain barrier function

**DOI:** 10.1101/2022.04.14.488066

**Authors:** Bárbara S. Casas, Gabriela Vitória, Catalina P. Prieto, Mariana Casas, Carlos Chacón, Markus Uhrig, Fernando Ezquer, Marcelo Ezquer, Stevens K. Rehen, Verónica Palma

## Abstract

Schizophrenia (SZ) is a complex neuropsychiatric disorder, affecting 1% of the world population. Long-standing clinical observations and molecular data have pointed out a possible vascular deficiency that could be acting synergistically with neuronal dysfunction in SZ.

As SZ is a neurodevelopmental disease, the use of human induced pluripotent stem cells (hiPSC) allows disease biology modeling retaining the patient’s unique genetic signature. Previously, we reported a VEGF-A signaling impairment in SZ-hiPSC derived neural lineages leading to a decreased angiogenesis. Here, we present a functional characterization of SZ-derived brain microvascular endothelial-like cells (BEC), the counterpart of the neurovascular crosstalk, revealing an intrinsically defective Blood-Brain Barrier (BBB) phenotype. Transcriptomic assessment of genes related to endothelial function among three control (Ctrl BEC) and five schizophrenia patients derived BEC (SZP BEC), revealed that SZP BEC have a distinctive expression pattern of angiogenic and BBB-associated genes. Functionally, SZP BEC showed a decreased angiogenic response *in vitro* and higher transpermeability than Ctrl BEC. Immunofluorescence staining revealed less expression and altered distribution of tight junction proteins in SZP BEC. Moreover, SZP BEC’s secretome reduced barrier capacities in the brain microvascular endothelial cell line HCMEC/D3 and in an in vivo permeability assay in mice. Overall, our results describe an intrinsic failure of SZP BEC for proper barrier function. These findings are consistent with the hypothesis that traces schizophrenia origins to brain development and BBB dysfunction.

## Introduction

Schizophrenia (SZ) is a complex chronic neuropsychiatric disorder, characterized by perturbations in thinking, perception, and behavior; along with brain connectivity deficiencies, neurotransmitter dysfunctions, and abnormal distribution of neurons in the prefrontal cortex [1]. SZ affects 1% of the population worldwide and, to date, it has no cure. Current pharmacological treatments are only partially efficacious with about 30% of patients describing little to no improvement after treatment [2].

It is now well accepted that SZ is a neurodevelopmental disease of multiple etiology, in which there is a basal genetic heterogeneity interacting with several environmental factors [3–6]. The aforementioned agents converge on the brain’s development, affecting pathways and processes that can trigger the disease that presents most commonly in the mid-to late 20’s [7–11]. Nevertheless, the mechanisms that can explain and predict the onset and evolution of the disease remain still largely unknown.

While most studies have focused on intrinsic neuronal deficiencies, long-standing clinical observations and increasing molecular evidence have linked SZ to vascular abnormalities, such as deficiencies in angiogenic factor levels, hypoperfusion, neuroinflammation, and blood-brain barrier (BBB) dysfunction [12–18].

Brain vascularization and BBB formation starts early during embryonic development by the recruitment of angioblasts and endothelial cells from the developing neural tube, forming the perineural vascular plexus. The newly generated blood vessels then migrate within the neural tube to form the new blood vessels from those pre-existent ones, a process known as angiogenesis [19]. Besides cell recruitment and angiogenesis, neural tissue induces the expression of specific transporters (such as GLUT-1) and reduces endothelial cell permeability. After cerebral angiogenesis starts, brain microvascular endothelial cells have diminished fenestrations and leakage and a higher expression of tight junction (TJ) proteins, such as Claudin-5 (CLN5) and Occludin (OCLN). These unique characteristics of the brain endothelium are collectively known as the BBB and confer them the ability to regulate molecular trafficking between blood and brain, assuring the delivery of oxygen and nutrients, and the removal of carbon dioxide and waste products from neural tissue [20].

Neuro-vascular coupling mechanisms during brain development assure the simultaneous generation of the neural network and blood vessels, and a proper BBB formation [21, 22]. In this context, it has been proposed that there is no BBB lineage but rather an initial phase of induction and establishment of the BBB phenotype, followed by a second phase that consists of the maintenance of BBB characteristics along adulthood [20].

SZ is among the most challenging conditions to study because of the complex nature of the human brain and the limitations of existing animal model systems in recapitulating all these uniquely human-specific traits, especially during neurodevelopment. With the recent advances in stem cell culture techniques, neurons derived from human induced pluripotent stem cells (hiPSCs), have shown to recapitulate neural development *in vitro* and serve as a model for various neuropsychiatric disorders, including SZ [23–25]. hiPSC conserve the genetic diversity of donors and have been used mainly for the modelling of neural-derived cells; being able to recapitulate classic hallmarks of the disease [24, 26–28]. Previously, we have shown that neural stem cells (NSC) derived from SZ-hiPSC induces an impaired angiogenesis, arguing in favor of a neurovascular dysregulation in this disease [29]. Based on the assumption that proper brain endothelial cells responsiveness in the brain is also critical for neural function, here we characterized endothelial cells derived from SZ-hiPSC and studied their functionality regarding two important processes of brain vasculature: angiogenesis and BBB formation. Our results reveal a possible intrinsic failure in the brain microvascular endothelial cells of SZ patients affecting proper angiogenesis and BBB function, that could contribute to an altered neurovascular crosstalk in SZ.

## Materials and Methods

### hiPSC differentiation to Brain Microvascular Endothelial-like cells

hiPSC were obtained from dermal fibroblasts reprogramming as described before [29]. We used three hiPSC lines obtained from healthy controls subjects (Ctrl hiPSC) and five from SZ patients (SZP hiPSC). We selected SZP hiPSC that came from patients with known familiar risk of SZ, assuring a high genetic component [30, 31]. Three SZP hiPSC lines were obtained from the Coriell Institute (SZP #1, SZP #2 and SZP #3) and two were reprogrammed at the D’Or Institute (SZP #4 and SZP #5). Regarding the Ctrl hiPSC we considered a similar age range than SZP hiPSC. One Ctrl hiPSC line was obtained from the Coriell Institute (Ctrl #1), while the other two hiPSC lines were reprogrammed at the D’Or Institute for Research and Education (Ctrl #2 and Ctrl #3). For detailed sample information refer to Supplementary Table 1. In all figures each donor cell line is identified in correlative numbers as described above, unless otherwise indicated.

For brain endothelium assessment hiPSC lines were derived to brain microvascular endothelial like cells (BEC) following previously published protocols [32, 33]. Briefly, hiPSC were dissociated with accutase and seeded at a density of 35×10^3^ cells/cm^2^ on Matrigel-coated dishes in mTeSR1 (STEMCELL Technologies) supplemented with 10 mM ROCKi Y-27632 10mM (Merck Millipore, Darmstadt, Germany) and medium was changed daily for 3 days. To initiate differentiation, cells were treated with 6 mM CHIR99021 (Merck Millipore) in DeSR1 media containing DMEM/F12 1% MEM-NEAA (Thermo Fisher Scientific), 0.5% GlutaMAX (Thermo Fisher Scientific), and 0.1 mM β- mercaptoethanol (Sigma-Aldrich). After 24 hours medium was changed to DeSR2: DeSR1 supplemented with B27 (Thermo Fischer Scientific). Medium was changed daily for 5 days. Next, medium was changed to hECSR1: hESFM (Thermo Fisher Scientific) supplemented with 20 ng/ml basic fibroblast growth factor (bFGF), 10 mM Retinoic Acid and B27. After 48 hours cells were dissociated with accutase and seeded at 1×10^6^ cells/cm^2^ in plates, Labtek chamber slides or transwell inserts (1.12 cm^2^ 0.4 µm Pore Polyester Membrane Insert, Corning) coated with Collagen IV (400 ng/ml)/Fibronectin (100 ng/ml). Next day medium was changed to Basal Medium: hESFM supplemented with B27. The 10^th^ day of differentiation was considered to be the first day of the BEC culture; these cells were used for all the analysis described below.

### Cell culture of HCMEC/D3 and HUVEC

The human brain microvascular endothelial cell line hCMEC/D3 (Merck Millipore) was seeded in T25 flasks coated with rat tail type I collagen (Merck Millipore), in EndoGro-MV medium (Merck Millipore) supplemented with 1 ng/ml of bFGF. Cells were used between passages 3 to 8.

HUVEC were obtained from full-term umbilical cords as previously described [34], seeded in dishes coated with 1% gelatin and cultured in primary cell medium composed by medium 199 (M199) supplemented with 10% NBCS, 10% FBS, 3.2 mM L-glutamine and 100 U/mL penicillin-streptomycin. The medium was changed every 2 days until confluence was reached, having a typical cobblestone appearance. All primary cultures of HUVEC were used between passages 2 to 6.

### Conditioned Medium (CM) Collection and Proteome Profiling

BEC were cultured in Basal Medium (hESFM supplemented with B27) for 24 hours. Medium was collected, centrifuged at 500xg for 5 minutes, aliquoted, fast frozen in liquid nitrogen and stored at -80ºC until use. For each analysis CM was pooled from 3 distinct differentiation rounds.

The presence of angiogenic factors in BEC CM was evaluated trough a Proteome Profiler Human Angiogenesis Array kit (ARY007, R&D Systems Inc., Minneapolis, MN, USA). For this, 650 µl of CM from each condition (3 Ctrl BEC and 5 SZP BEC) were assayed. Signal was detected by enhanced chemiluminescence with an UVITEC 4.7 Image System (Cambridge, UK) and intensity was quantified by densitometry using the software ImageJ (NIH, USA). The pixel intensity of each factor (in duplicate) was normalized to that of the three internal controls provided by the assay. Each assay was performed in duplicate.

### Immunofluorescence

Cells seeded on Labtek or transwell were washed with PBS, fixed with cold methanol, permeated with 0.01% Triton (Sigma, St. Louis, MO, USA) solution, blocked with 3% BSA and incubated with the following primary antibodies: rabbit anti-CD31 (1:50, PA516301 Invitrogen), mouse anti-GLUT1 (1:100, MA5-11315 Invitrogen), mouse anti-OCLN (1:50, sc-133256 Santa Cruz Biotechnologies), mouse anti-CLN5 (1:20, 35-2500 Invitrogen), mouse anti-ZO1 (1:50, 33-9100 Invitrogen). Subsequently, cells were incubated with secondary antibodies: Alexa-555 anti-rabbit (1:500, A21429 Invitrogen) and Alexa-488 anti-mouse (1:500, A11029 Invitrogen), respectively. Phalloidin and DAPI staining were used for actin and nuclei visualization, respectively. Images were acquired with a confocal microscope (Carl Zeiss 710) using the image acquisition program Zen (Carl Zeiss). Images were processed and analyzed using ImageJ software (NIH).

### Western Blot

Ctrl and SZP BEC, HUVEC and HCMEC/D3 were harvested in monolayers. Homogenates were suspended in SDS lysis buffer with added protease and phosphatase inhibitors (Catalog # 88265; Thermo Scientific, Waltham, MA, USA) at 4 °C. Equal amounts of protein were separated by 8–12% SDS-PAGE, followed by Western blotting using anti-ZO-1 (225-130 kDa), anti-Claudin-5 (18 kDa), anti-eNOS (130 kDa, sc C-654 Santa Cruz Biotechnologies) anti-FLK-1 (150 kDa, sc6251 Santa Cruz Biotechnologies) and anti-occludin (60-82 kDa) antibodies incubated overnight at 4 °C. The monoclonal mouse anti-β-actin (42 kDa, Sigma-Aldrich) antibody was used as a loading control. Membranes were washed in Tris-buffered saline (TBS) with 0.1% Tween, and incubated (2 h, 22 °C) in TBS/0.1% Tween containing horseradish peroxidase-conjugated goat anti-mouse secondary antibodies. Bands were visualized using enhanced chemiluminescence (ECL; Amersham Biosciences, Little Chalfont, UK), and quantified by densitometry using ImageJ (NIH, USA).

### Gene expression analysis

Total RNA was obtained from all BEC lines (3 Ctrl BEC and 5 SZP BEC) by phenol-chloroform extraction using RNAsolv (Omega Bio-Tek, Norcross, GA, USA). 1 μg of RNA was treated with DNase I (Invitrogen) and then cDNA was synthesized using M-MLV reverse transcription kit (Promega, Madison, WI, USA). Relative expression was assessed by qPCR with SyberGreen II mix (Agilent Technologies Thermocycler, Santa Clara, CA, USA) using specifically designed primers as indicated in Supplementary Table 2. Data was analyzed by calculating the expression fold change via 2^- ΔΔCt^, and gene expression was normalized to that of three reference genes (*GAPDH, B2M,* and *18S*). For each analysis, cDNA from BEC of two distinct differentiations procedures was used.

### Tube formation assay

To evaluate angiogenic capacities of BEC, we performed a tube formation assay as previously described [34]. Briefly, cells (50.000/well) were seeded onto solid growth factor-reduced Matrigel (BD Biosciences, San Jose, CA, USA) in 96-well plates with the following stimuli: Basal Medium (hESFM supplemented with B27), Basal Medium plus 50 ng/ml human recombinant VEGFA (PHC9391 Gibco, Frederick, MD, USA) or Basal Medium plus 100 ng/ml recombinant human NTN1 (6419-N1, R&D Systems, Minneapolis, MI, USA). Photographs were taken after 4 h of incubation. Tubes and sprouts were quantified using Angiogenesis Analyzer for ImageJ [35]. Example of segmentation of images on Figure 2a can be found in Supplementary Figure 2d.

To evaluate the angiogenic potential of BEC CM a tube formation assay was performed using the HCMEC/D3 line (SCC066 Merck Millipore, Darmstadt, Germany), as described above, with the following stimuli: EGM as positive control, Basal Medium as negative control, CM from Ctrl BEC, CM from SZP BEC and CM from HUVEC. All assays were performed in triplicate.

### Measurement of trans-endothelial electrical resistance (TEER)

BEC were cultured in transwell inserts (1.12 cm^2^ 0.4 µm Pore Polyester Membrane Insert, Corning) previously coated with Collagen IV /Fibronectin at a density of 9×10^5^ cells/cm^2^. TEER was measured with an epithelial Volt/Ohm meter (EVOM2, World Precision Instruments, Sarasota, FL) once a day for three days. For each BEC line, measurements were performed on 4 distinct differentiation procedures.

For evaluation of BEC CM, HCMEC/D3 were cultured in transwell inserts (1.12 cm^2^ 0.4 µm Pore Polyester Membrane Insert, Corning) previously coated with rat tail type I collagen, at a density of 25×10^3^ cells/cm^2^ and cultured in EndoGro medium (Merck Millipore, Darmstadt, Germany). TEER was measured daily and experiments were performed when it reaches over 30 Ωcm^2^ (5-7 days after seeding). CM was added to the apical chamber and Basal Medium to the basolateral chamber. The TEER was measured 1 h after medium change for monolayer stabilization (Time 0) and 24 h afterwards. To evaluate the effect of BEC CM on BBB permeability we calculated the percentage of change of the TEER between 24 h and the basal condition (time 0). For each CM, experiment was repeated twice.

### *In vitro* permeability assay

For permeability experiments, BEC were cultured in transwell inserts (1.12 cm^2^ 0.4 µm Pore Polyester Membrane Insert) previously coated with Collagen IV /Fibronectin at a density of 9×10^5^ cells/cm^2^. Cells were washed twice with PBS and then the medium was replaced with hESFM (ThermoFischer). Fluorescein isothiocyanate (FITC)-dextran 70 kDa solution was added into the apical chamber at a 1 µM final concentration. 150 µL of medium in the basolateral compartment were collected after 60, 80, 100 and 120 min and stored at −20 °C, each time the volume was replaced with hESFM. The fluorescence in the collected media was measured using a Nano Quant Infinite 200 ProNano (Tecan, Männedorf, Switzerland) at 488/530 nm. The background values (hESFM reading) were subtracted from every measurement. For each assay, BEC from two distinct differentiation procedures were used.

### *In vivo* vascular permeability assay

To evaluate the in vivo effects of Ctrl and SZP BEC CM on vascular permeability, we performed a Miles assay as previously published [36, 37]. In order to reduce the number of animals used in this study, Ctrl and SZP BEC CM were pooled from 3 different donors each.

8-10 weeks old male Balb/c mice were obtained from the Faculty of Medicine’s Animal Facility, Universidad de Chile. Room temperature was maintained at 23°C, with a 12:12 h light-dark cycle. Mice were fed *ad libitum*. All procedures were approved and supervised by the Bioethics Committee of the Faculty of Medicine, Universidad de Chile (protocol CBA-1103-FMUCH).

For Miles assay mice were intraperitoneally (i.p.) injected with a solution of Pyrilamine maleate (4 mg/mL) using 10 µL/gram of mouse. After 20 min, mice’s tails were wrapped with a warm gel pack (available in commerce), for 1-3 min to increase vasodilatation. Then, 100 µL of Evan Blue solution (1% w/v) were intravenously (i.v.) injected in the tail. After 30 min, mice were anesthetized as described and 40 µL of each BEC CM, Basal Medium or VEGFA were locally injected intradermally (i.d.) in 3 different points in the flanks of the mice (BEC CM Media was injected in one side of the mouse and Basal Medium and/or VEGFA on the other side as an internal control) and let recover in their home cages. Mice were euthanized 35 min after the procedure to recover the skin regions surrounding i.d. injections. Skin samples were stored at -20°C. Dye was extracted with formamide and quantified by absorbance measuring at 620 nm with a reference reading of 740 nm using a spectrophotometer. Evans blue leakage was calculated as the absorbance of samples treated with BEC CM normalized against absorbance of samples injected with Basal Medium in the same mouse.

### Statistical Analysis

For all experiments three biological replicates were considered for the Ctrl group (Ctrl #1-3) and five for the SZ group (SZP #1-5), unless otherwise indicated in figure legends. All statistical analyses were performed using Graphpad Prism 9.0 (GraphPad Software Inc). Normality was assessed with D’Agostino-Pearson test. Nested t-test was used for comparison between two groups. Two-way ANOVA was used for multivariable approximations. Statistical significance was set at p < 0.05.

Past v4.02 (Ø, Hammer and D.A.T. Harper) software was used for Principal Component Analysis.

## Results

Nowadays, efficient hiPSC *in vitro* modeling permits the understanding of the molecular mechanisms of BBB development and its regulation. Our hiPSC-derived brain microvascular endothelial like cells (BEC) present a clear endothelial morphology (Supp Fig 1a, b) and expressed classical endothelial markers such as VEGF receptors (*FLT-1* and *KDR*), *PCAM, CDH5, vWF* and eNOS, as well as genes coding for specific BBB markers such as tight junction proteins *OCLN, ZO-1, CLN5*, and transporters *P-gP* and *GLUT-1* (Supp Fig 1b-f). Expression levels of these genes have similar values compared to the brain microvascular endothelial cell line hCMEC/D3 and primary endothelial cells HUVEC ones(Supp Fig 1 d-f). BEC also exhibit high TEER values, similar to those encountered *in vivo* (Supp Fig 1h).

### SZP BEC have a distinctive expression pattern of function-associated genes

Having established an efficient BEC differentiation protocol, we derived cultures from three healthy control donors (Ctrl BEC #1-3) and five patients with schizophrenia (SZP BEC #1-5). We assessed transcriptional levels of genes associated to main brain endothelial functions: angiogenesis and BBB. We measured classic angiogenesis-related genes, such as VEGF receptors, FLT-1 and KDR, VEGFA co-receptors *NRP1* and *NRP2, HIF1a* and *uPA* (Figure 1a), and BBB-related genes, such as *GLUT1, P-gP, CLN5, OCLN, ZO-1* (Figure 1b). Most of the genes analyzed presented variability in their expression levels between cell donors; nevertheless, this variability was lower within the Ctrl group when compared to the SZP group. When comparing mRNA levels of these genes between Ctrl and SZP BEC group, we found that three genes showed significant differences.

mRNA levels of *HIF1a*, whose pathway has been proposed to be upregulated in SZ [38], was increased in SZP BEC 3.4 times compared to the level in Ctrl BEC (Figure 1a).

**Figure 1.**
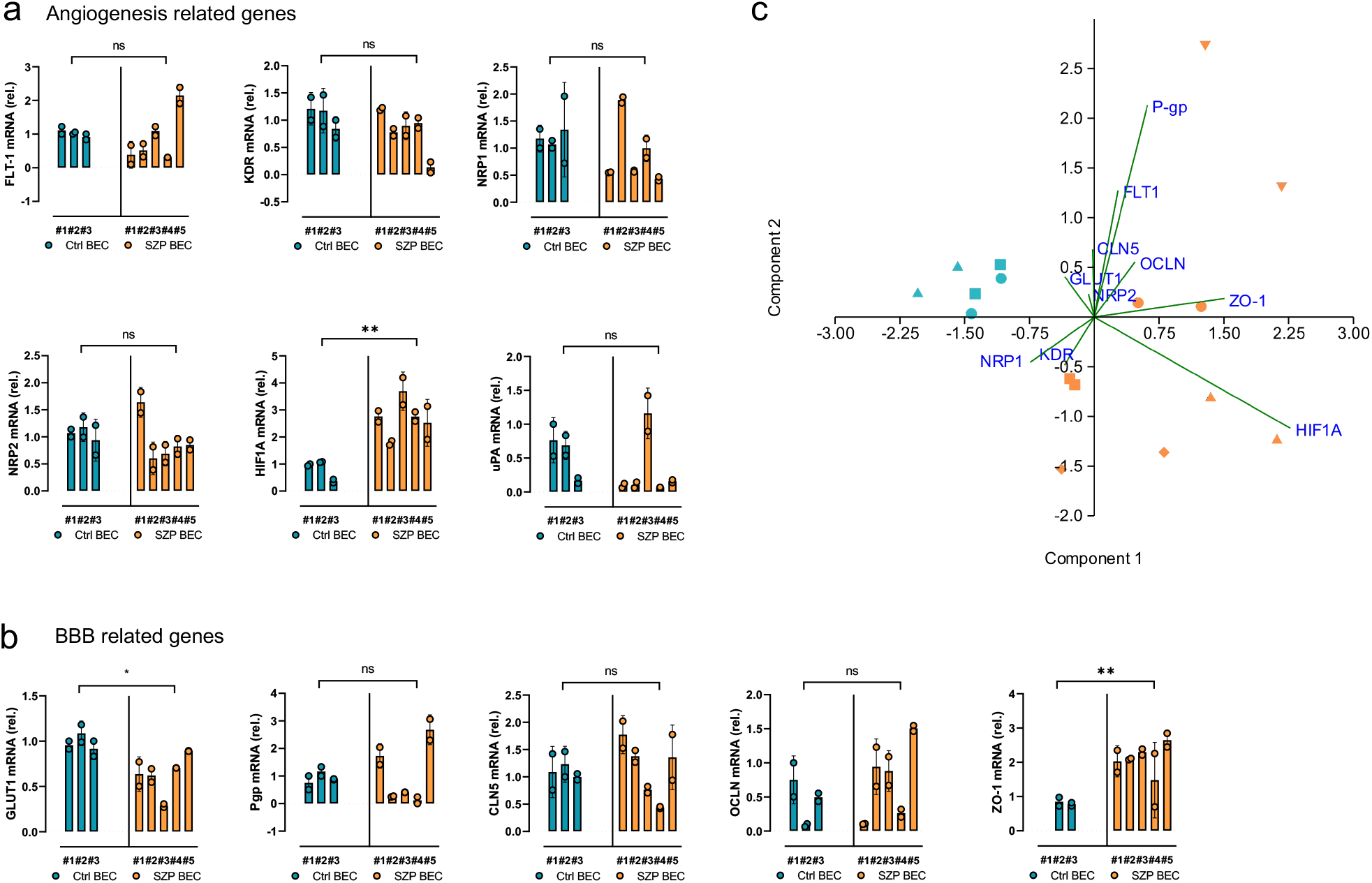
SZP BEC have an altered gene expression pattern of angiogenic and BBB-related genes. Eight hiPSC, three Ctrl and five SZP, were differentiated to BEC. **a-b** RNA was extracted from two independent differentiations rounds for each donor and mRNA levels of angiogenic (a) and BBB-related genes (b) were assessed by RT-qPCR; *B2M* was used as housekeeping gene. Data are expressed as mean ± SD, with * p< 0.05 and ** p≤0.01 according to Nested t test. c Graph shows values of PC1 and PC2 of PCA performed on (a-b).

Regarding BBB-related genes, mRNA levels of GLUT-1 levels were lower in SZP BEC, 0.64 times the levels found in Ctrl BEC (Figure 1b). ZO-1 was 2.6 times higher in SZP BEC, in comparison to the levels measured in Ctrl BEC (Figure 1b).

Since SZ has been described as a polygenic disease, where multiple gene variants may converge in similar molecular pathways or gene modules, we aimed to evaluate if the high expression variability among SZP BEC could result in a distinctive SZP pattern. Therefore, we performed a Principal Component Analysis (PCA) of the evaluated gene levels and confirmed that the first component allows us to discriminate between the Ctrl and the SZP group (Figure 1c), which may be a product of a distinctive expression pattern in these cells. Among the genes with higher weight of this first component we found: *HIF1A, ZO-1, NRP1* and *PgP*. Interestingly, *CLN5* had a minimum weight in differentiating Ctrl and SZP BEC groups (Supplementary Figure 1g).

### SZP BEC have a lower response to angiogenic stimuli

In order to evaluate the intrinsic angiogenic capacity of BEC, we performed a tube formation assay. No differences were found between Ctrl and SZP BEC when incubating in basal culture medium (Figure 2a, Supplementary Figure 2).

**Figure 2.**
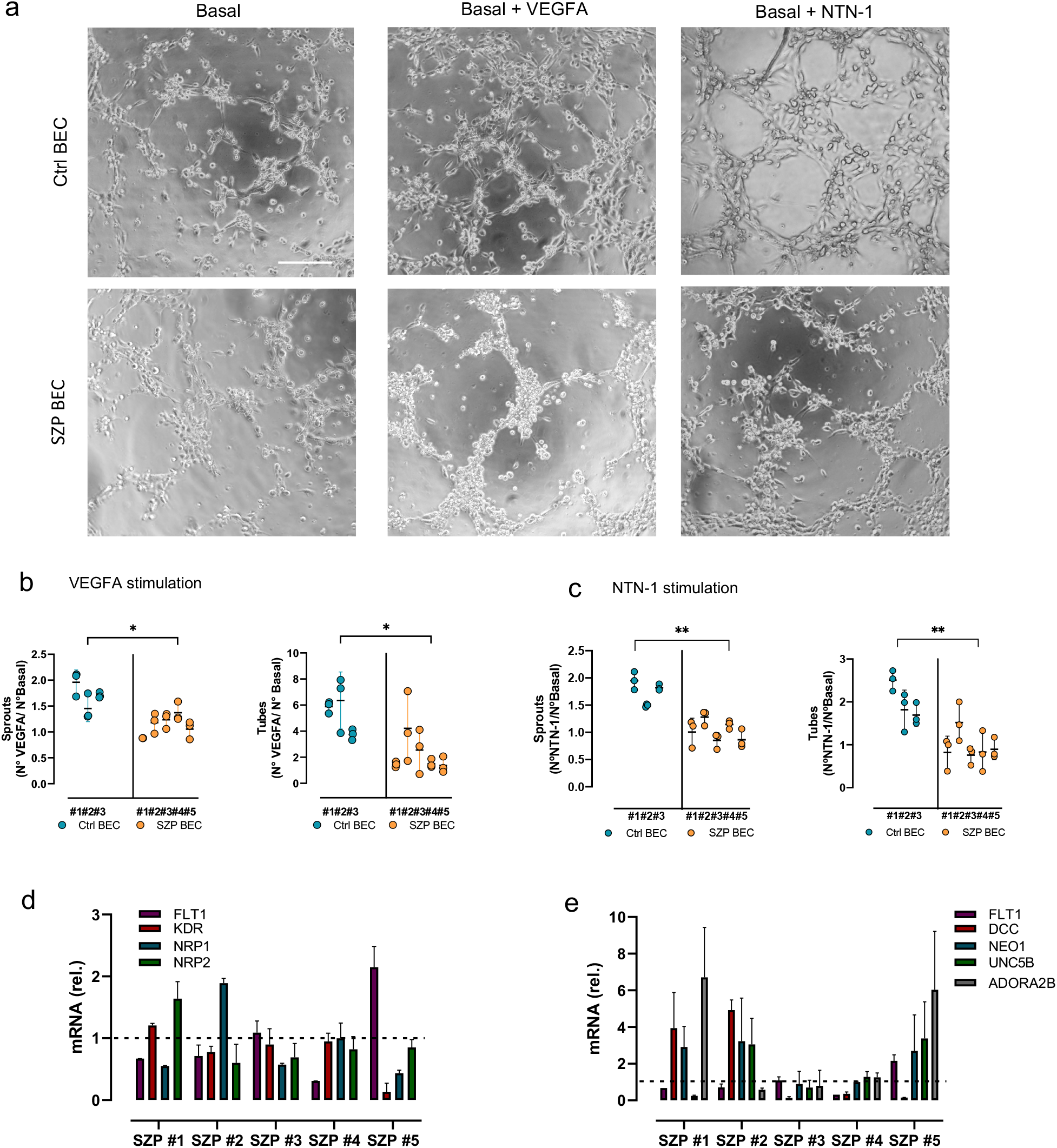
SZP BEC present a lowered response to an angiogenic stimuli. **a.** Representative images of tube formation assay when incubating Ctrl and SZP BEC with basal culture medium; supplemented with VEGFA 50ng/ml or with NTN-1 100ng/ml, scale bar= 30 µm. For image segmentation refer to Supplementary Figure 2d. **b-c** Quantification of fold increase in average number of sprouts and tubes generated after stimulation with 50ng/ml VEGFA (b) or 100 ng/ml NTN-1 (c) respect to the basal culture medium. VEGFA and NTN-1 stimulation was performed in triplicate for each cell donor. * p < 0.05, ** p <0.01, according to Nested t-test. **d.** Quantification of mRNA levels of *FLT-1, KDR, NRP-1* and *NRP2*, for each SZP BEC line assayed in (a). Levels are normalized against B2M (housekeeping) and graphed as fold increase of average Ctrl BEC values (represented as segmented line = 1). Data are shown as mean ± SD.

Since VEGFA expression has shown to be downregulated in SZ and in SZP-derived NSC, we next evaluated the angiogenic response of BEC to VEGFA stimulation [29, 39–42].

VEGFA stimulation of Ctrl BEC increased *in vitro* angiogenesis, whereas no qualitative difference was observed when stimulating SZP BEC (Figure 2a, Supplementary Figure 2d). When comparing fold increase between basal and VEGFA stimulated conditions, we observed that Ctrl BEC formed sprout and tubes in a higher proportion than SZP BEC (Figure 2b, c). VEGFA increased Ctrl BEC sprout formation by 70% whereas in SZP BEC the increase was only 16% (Figure 2b). Ctrl BEC formed 5.3 times more tubes in response to VEGFA. In turn, in SZP BEC tubes increased only 2.2 times (Figure 2b).

The non-canonical angiogenic molecule, NTN-1, is relevant for both angiogenesis and axonal guidance [43–45]. NTN-1 induced angiogenesis in Ctrl BEC, whereas no significant change was observed in SZP BEC after stimulation (Figure 2a). When comparing the fold increase in sprout and tube formation after NTN-1 simulation, we observed a 70% increase in sprout formation in Ctrl BEC and no increase in SZP BEC. Ctrl BEC generated 2 times more tubes in response to NTN-1, but the ligand had no effect in SZP BEC (Figure 2c).

As observed in Figure 1, no significant alteration was observed in the expression of any of the VEGFA receptors or co-receptors when comparing between Ctrl and SZP BEC groups. Nevertheless, when assessing individual transcriptional values of all receptors in each SZP BEC line, we observed that the overall pattern of expression differs from the Ctrl BEC (represented as an average value=1) (Figure 2d). Similarly, NTN-1 receptors show great variability of expression between cell donors (Figure 2e).

### SZP BEC exhibit higher permeability than Ctrl BEC

As stated before, one of the main functions of the brain endothelium is the structuration and maintenance of the BBB, which results in a decreased permeability of solutes across the endothelial cells. To asses this function we cultured Ctrl and SZP BEC in transwell inserts and measured TEER of monolayers within 48 hours of cell culture. hiPSC-derived BEC generate high TEER values, as reported before [32]. TEER had a maximum at day 1 of cell culturing with an average value of 1179 ± 504 Ω (measurements for each cell line can be observed in Supplementary Figure 3a).

When comparing Ctrl and SZP BEC groups we observed that during an equivalent time course of measurements, SZP BEC had lower TEER values than Ctrl SZP BEC (Figure 3a). In addition, we measured permeability to dextran within a two-hour period. We observed that permeability was significantly higher in SZP BEC than in Ctrl BEC after 60 min of incubation, permeating 1.8 times more dextran than Ctrl BEC (Figure 3b). After this time the mean permeability tended to be higher in the SZP group, but had no longer statistical significance (Supplementary Figure 3b-c).

**Figure 3.**
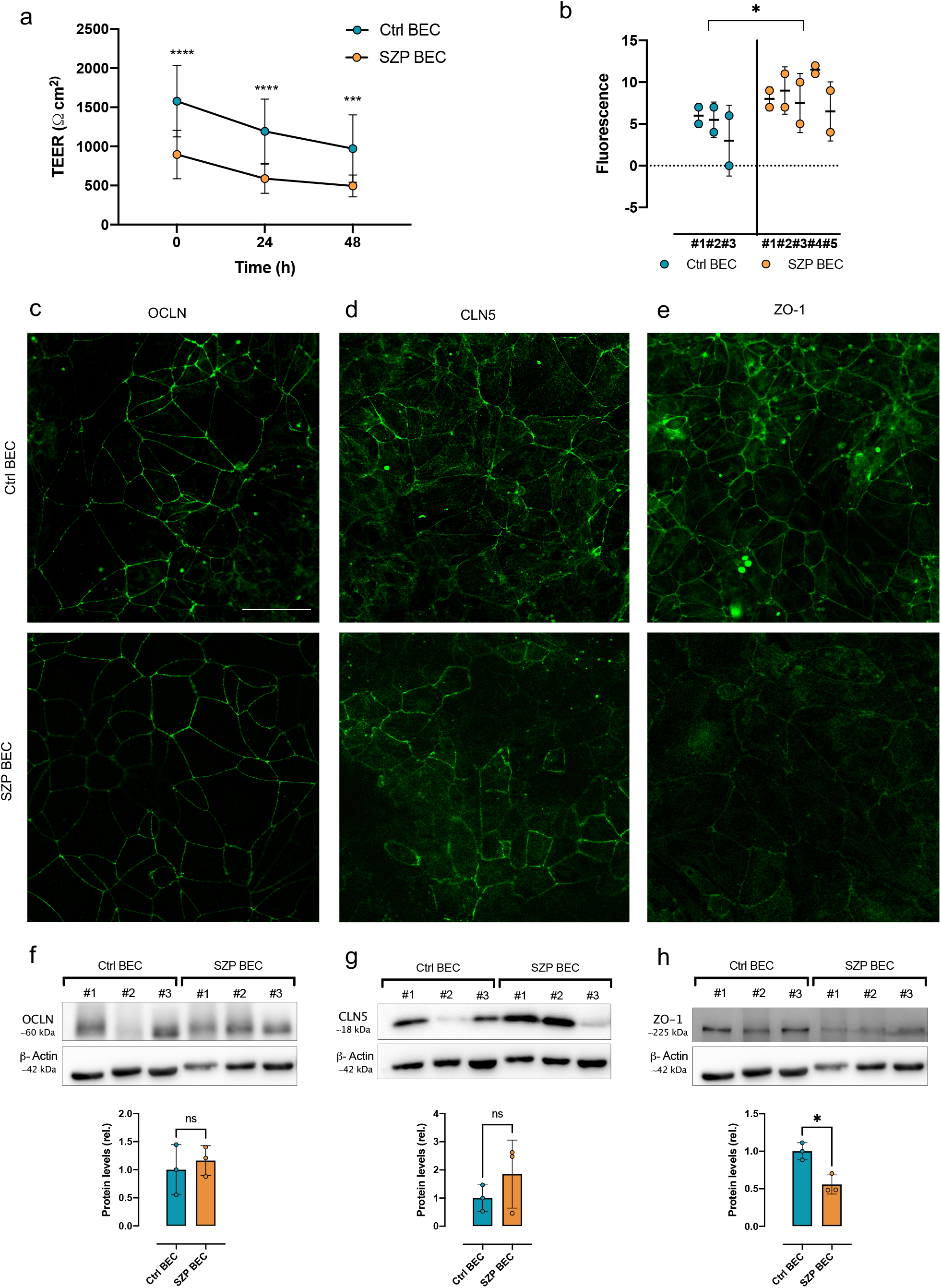
SZP BEC exhibit an increased transpermeability. **a** TEER was measured on the first day of BEC differentiation (time=0 h) and after 24 and 48 h of culture. Data are shown as mean ± SD and include all biological replicates of Ctrl and SZP groups. *** p< 0.001, **** p< 0.0001 according to two-way ANOVA. **b** Dextran (fluorescence) permeability was assessed in Ctrl and SZP BEC. Data are shown as mean ± SD for each donor, with * p< 0.05 according to Nested t-test. **c-d** Representative immunofluorescence of OCLN (c), CLN5 (d) and ZO-1 (e) in Ctrl and SZP BEC; scale bar= 50 µm. **f-h** Total protein was extracted from whole cell lysates from two independent differentiations for 3 Ctrl and 3 SZP donors (indicated with correlative numbers). Western blot was performed on pooled samples of each donor to assess the expression of OCLN (f), CLN5 (g) and ZO-1 (h). Graphs show the mean expression level of each protein normalized against β-Actin and Ctrl BEC samples. Data are shown as mean ± SD of two independent blots, with * p<0.05 according to t-test.

As expected for the high TEER values obtained, both Ctrl and SZP BEC express TJ proteins at the cell membrane (Figure 3c-d). The protein expression and distribution of OCLN seams equivalent between Ctrl and SZP BEC (3c). Nevertheless, when assessing CLN5 expression and distribution, we observe a fair and continuous staining at the cell membrane of Ctrl BEC, whereas in SZP BEC we found regions of absent or noncontinuous staining at the cell membrane and a more cytoplasmic distribution (Figure 3d, Supplementary Figure 3d). In SZP BEC, ZO-1 seems to have a weaker expression with a more cytoplasmic distribution compared with the high and characteristic cell membrane restricted distribution of ZO-1 in Ctrl BEC (Figure 3e, Supplementary Figure 3e).

In line with our transcriptional level results, we did not find any significant difference in OCLN and CLN5 protein expression levels (Figure 3f-g). On the other hand, contrary to the increased mRNA levels found for ZO-1 in SZP BEC, we observed that ZO-1 protein levels were 45% lower in SZP group compared to Ctrl BEC (Figure 3h). This differences in TJ protein expression and distribution could explain, in part, the decreased TEER values and higher permeability of SZP BEC.

### Angiogenic profiling of BEC secretome

We have previously shown that hiPSC-derived SZP NSC induce an impaired angiogenesis and present an altered angiogenic secretome, with a decreased expression of pro-angiogenic molecules, which could be affecting paracrine angiogenic signaling in the neurovascular niche [29]. We aimed therefore to analyze if there was also an imbalance in angiogenic molecule secretion in SZP BEC, whicht could be contributing to the decreased angiogenic signaling observed in NSC. For that, we collected the conditioned medium (CM) of each cell line.

Angiogenic profiling of SZP and Ctrl BEC CM revealed a similar level for most detected proteins (Figure 4a). Two of the 55 factors assessed showed a significant difference between Ctrl and SZP groups: MMP9 and uPA (Figure 4b-c). MMP9, a matrix metalloproteinase that is important for both angiogenesis and synaptic plasticity [46], was found to be 2.9 times higher in SZP BEC CM than in Ctrl BEC ones (Figure 4b). Interestingly, MMP9 upregulation has been linked to BBB disruption [47]. In turn, uPA was 70% lower in SZP BEC compared to the Ctrl group (Figure 4c). Concordantly, uPA was also found to be reduced in SZP NSC [29].

**Figure 4.**
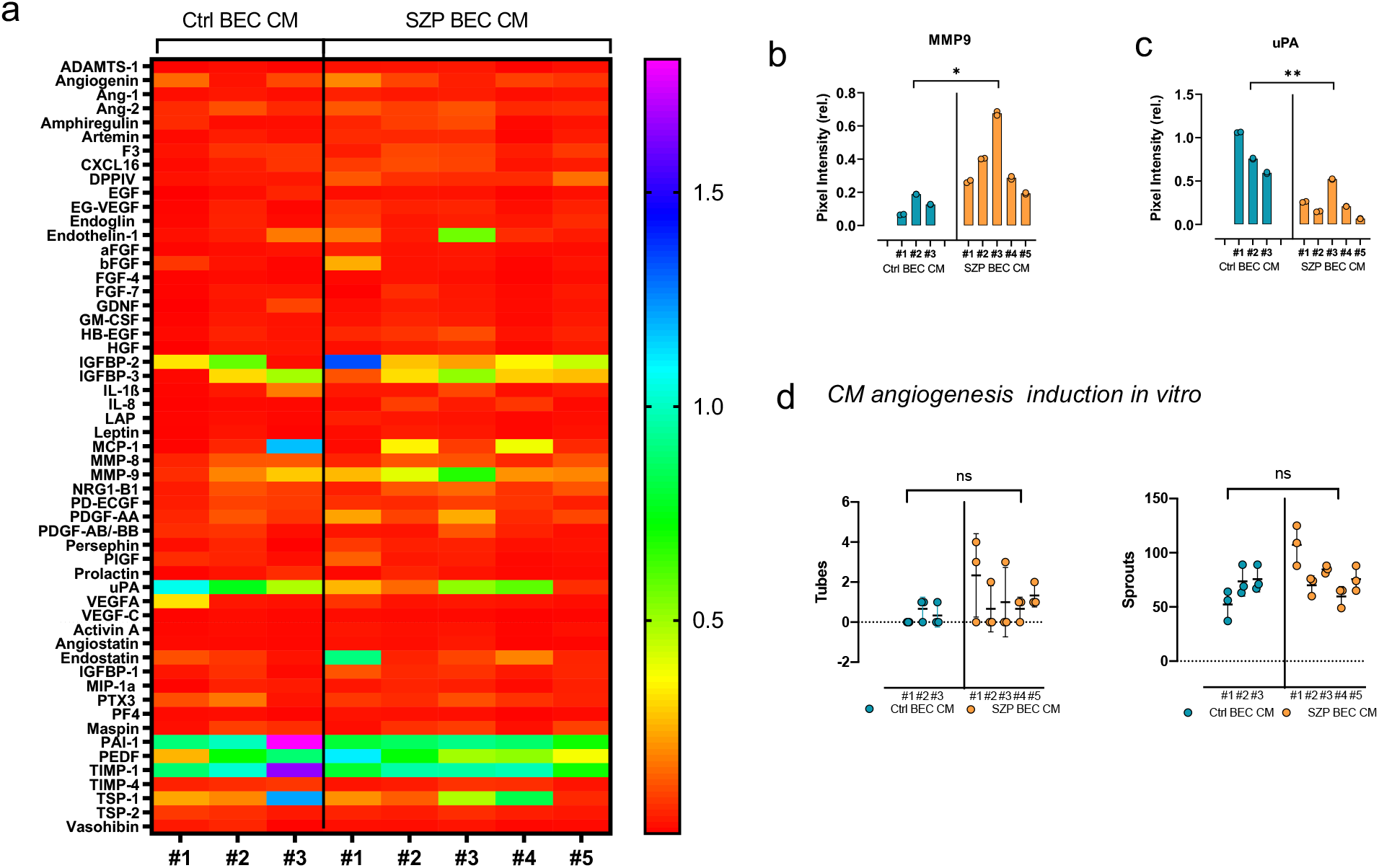
SZP BEC CM present a distinct angiogenic profile. **a** Heatmap of angiogenic factors detected in BEC CM. Data are shown as mean level of each factor in CM from each cell donor respect to an internal control of the assay. Color key at right. **b-c** Graph shows levels of MMP9 (b) and uPA (c) detected in Ctrl and SZP BEC CM. Data are expressed as mean ± SD, with * p< 0.05 and ** p≤0.01 according to Nested t test. **d** Quantification of average number of sprouts and tubes generated after stimulation of HCMEC/D3 with either Ctrl or SZP BEC CM. Stimulation was performed in triplicate for each cell donor CM.

To evaluate the possible distinct autocrine effect of Ctrl and SZP CM over angiogenesis, we performed a tube formation assay on the brain microvascular endothelial cell line (HCMEC/D3). We found no differences in angiogenesis induction, neither in sprout nor in tube formation, between Ctrl and SZP CM groups (Figure 4d). Moreover, angiogenesis induction was similar to the one induced by HUVEC CM (Supplementary Figure 4).

### SZP BEC CM increases vascular permeability

Since endothelium-derived signals can have an autocrine effect impacting BBB integrity [48], we next evaluated permeability alterations caused by Ctrl and SZP BEC CM both *in vitro* and *in vivo*.

HCMEC/D3 were seeded on transwell inserts and treated with either Ctrl or SZP BEC CM. Ctrl BEC CM increased HCMEC/D3 TEER by 20%, whereas SZP BEC CM reduced TEER about 13% (Figure 5a). When evaluating the presence of TJ protein, we observed a reduction of CLN5 and OCLN staining in HCMEC/D3 treated with SZP BEC CM (Figure 5b). SZP BEC CM reduced CLN5 expression in 19% and OCLN expression in 27% with respect to the Ctrl BEC CM (Figure 5b-d). In an intriguing manner, ZO-1 had a 45% increased expression in cells treated with SZP BEC CM (Figure 5e), nevertheless the distribution of ZO-1 in those cells appeared misallocated, with a higher cytoplasmic staining (Figure 5b).

**Figure 5.**
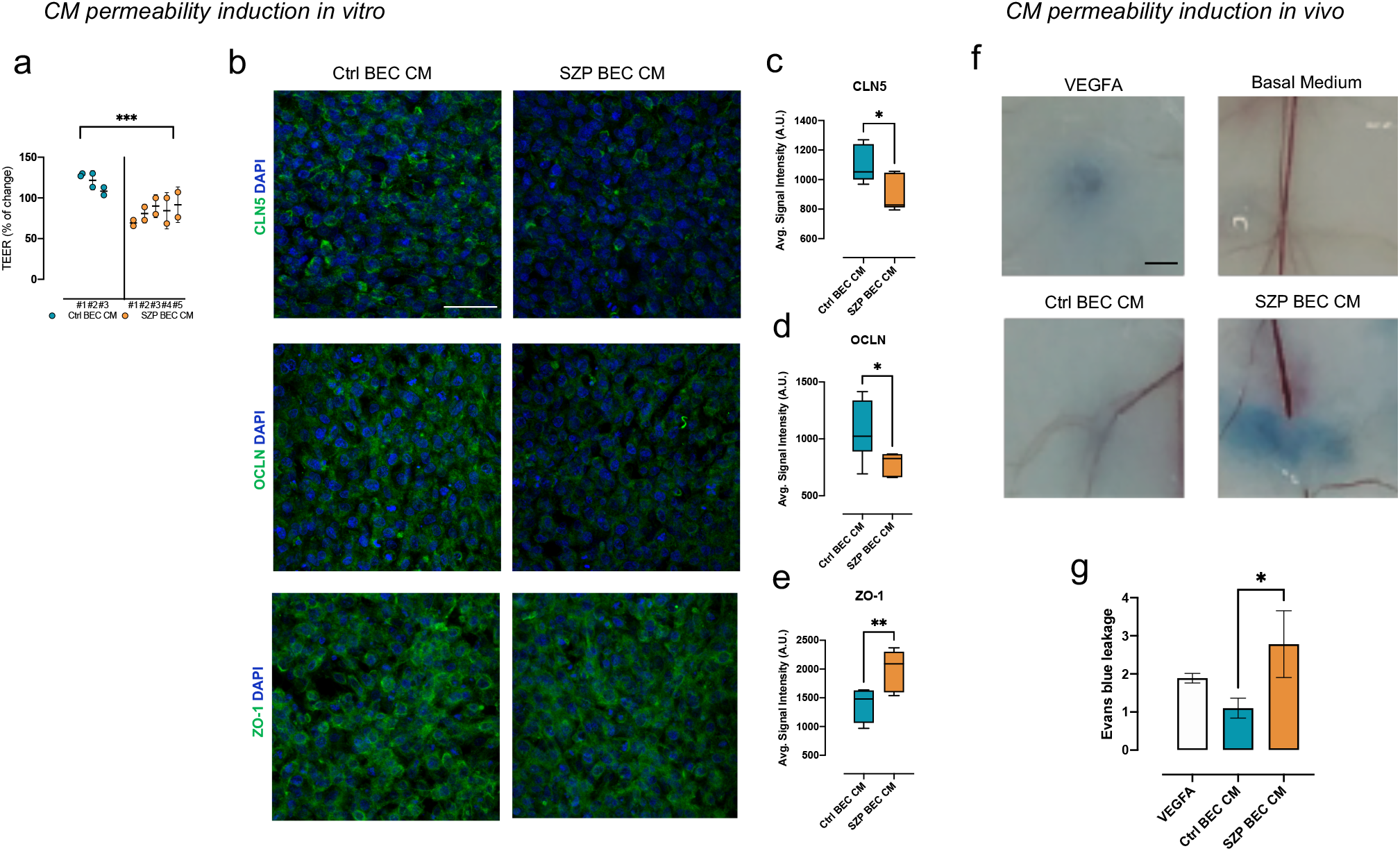
SZP BEC CM increases vascular permeability. **a** TEER was measured after 24 h incubation of HCMEC/D3 with BEC CM of each cell donor, identified in correlative numbers (Ctrl BEC CM #1-3 and SZP BEC CM #1-5). Data are shown as mean ± SD, *** p< 0.001 according to Nested t-test. **B** Representative immunofluorescence staining of CLN5, OCLN and ZO-1 of HCMEC/D3 transwells incubated with Ctrl or SZ BEC CM. Images show average intensity projections (11.6 µm depth for CLN5 and ZO-1; 4.7 µm depth for OCLN); scale bar= 50µm. **c-e** Quantification of average signal intensity of images on (b). Data includes all biological replicates of Ctrl and SZP group; * p< 0.05 and ** p≤0.01, according to t-test. **f** Representative photographs show Evans blue leakage in mouse skin injected either with 100 ng VEGFA, Basal Medium (hESFM + B27), Ctrl BEC CM or SZP BEC CM. Scale bar= 2 mm. **g** Quantification of Evans blue leakage in mouse skin calculated as absorbance at 620 nm with a reference reading at 740 nm and normalized against readings of Basal Medium injections in the same mouse. N=6 mice, * p<0.05 according to ANOVA.

Finally, we performed a Miles assay using pooled CM from Ctrl and SZP BEC on mice skin, using VEGFA as positive control (Figure 5f). SZP BEC CM increased vascular permeability respect to Basal Media and Ctrl BEC CM, as can be identified by an increase in Evans blue leakage around the injection site (Figure 5f). Quantification of the dye in the collected skin surrounding the injection site indicated that SZP BEC CM increases the leakage of Evans Blue 2.7 times compared to Ctrl BEC CM (Figure 5g). These results indicate that SZP BEC CM possesses a distinctive secretome that can alter vascular permeability *in vivo*.

## Discussion

In the present work we show that BEC derived from SZ patients present an intrinsic failure in two main processes important for proper brain vascularization: angiogenesis and BBB function.

We evaluated the expression of classical angiogenic and BBB-related genes. Most of the assessed genes did not show any significant difference between Ctrl and SZP groups. Similarly to what has been described in human post-mortem brains, we observe an increased gene expression variability in SZP BEC compared to Ctrl BEC [49]. Nevertheless, when analyzing the expression of all these genes by a PCA we could segregate clearly the Ctrl from the SZP BEC group, indicating that, although there is no significant association of many of endothelial related genes with SZ, the overall pattern of expression is characteristic for the SZP group. In addition to this, we found that three over five of genes with higher weight were related to BBB function. Moreover, we observed a reduced angiogenic response when stimulating SZP BEC with either VEGFA or NTN-1 and a disruption in barrier function. These observations are consistent with the occurrence of biological convergence in SZ [24], i.e., genetic variability among patients could be converging in the same cellular function deficiency leading to a BBB dysfunction.

Among the differentially regulated genes we found an upregulation of *HIF1A* and *ZO-1* and a downregulation of *GLUT-1*. HIF1A is the most important mediator of the hypoxic response, regulating several angiogenic and metabolic genes, including *VEGFA* and *GLUT-1* [50]. Metagenomic analysis have shown an upregulation of many transcriptional targets of HIF and other hypoxia related genes in SZ [51]. This paradox regarding an upregulation of HIF and a downregulation of VEGFA reported in SZ, remains elusive and more information regarding HIF1A pathway regulation in SZ is needed, especially regarding protein expression and stability.

In SZP BEC ZO-1 failed to localize properly at the cell membrane, revealing an altered expression pattern with a more cytoplasmic distribution. ZO-1 is a junctional adaptor protein located at the inside of the cell membrane This protein has been shown to be crucial, not only for barrier formation, but also for CDH5-mediated adherens junctions, for cell-cell tension and for angiogenesis [52]. Noteworthy, SNPs associated to SZ have been found in PDZ motifs (ZO-1 has a PDZ motif) for two other genes, *PICK1* and *PDLIM5* [53, 54]. mRNA levels of those genes are elevated in SZ and the presence of SNPs is associated to functional deficits [53, 54]. We describe increased mRNA levels of *ZO-1* in SZP BEC compared to Ctrl BEC. It has been proposed that ZO-1 transcriptional expression could be regulated by a mechanism that includes both HIF1A and the schizophrenia-associated transcriptional factor ZEB1 [55], a matter that should be further investigated. Interestingly this transcript increase did not traduce in an increment of protein levels, rather we found a significant decrease in ZO-1 protein expression in the SZP BEC group. A deeper knowledge of ZO-1 expression and genetic alterations is needed in order to decipher its functional implications in the SZ pathology. In addition, we also observe disruption in CLN5 localization at the cell membrane in SZP BEC. These two observations were consistent with the decreased TEER and the higher permeability observed in SZP BEC when compared to Ctrl BEC.

Our results are in line with recent observations indicating a BBB disruption in SZ hiPSC-derived BEC [56–58]. These authors showed that BEC derived from a SZ patients with the 22q11.2 deletion syndrome (22q11DS) present lowered TEER values, increased permeability and decreased localization of ZO-1 and OCLN at the cell membrane [56, 58]. The presence of 22q11DS in SZ is often traduced in CLN5 downregulation which itself increases BBB permeability [59, 60]. Since the SZP cell lines used in this study do not have the 22q11.2 deletion syndrome, our results indicate that BBB disruption maybe a more general feature of the SZ endothelium, arising from different genetic landscapes.

Interestingly, when treating the BBB cell line (HCMEC/D3) with SZP BEC CM, a BBB-disruption, but no alteration in angiogenesis, was observed, indicating a possible dysregulation in autocrine signaling for BBB-maintenance in SZP BEC [61]. SZP BEC CM also produced a significant increase in vascular leakage in vivo, corroborating our *in vitro* assessment. Treatment with SZP BEC CM decreased CLN5 and OCLN immunostaining on HCMEC/D3 cultures, consistent with a decrease in the TEER after exposure to this secretome. We found an increase in MMP9 in SZP BEC CM compared to Ctrl BEC CM. It has been reported that treatment with MMP9 decreases TJ protein immunostaining and that MMP9 signaling impairs BBB function [47, 62]. Moreover increased MMP9 in plasma of patients with SZ has been described, along with SZ associated polymorphisms [63–68]. Hence, the herein described MMP9 increased activity could be one key factor associated with BBB disruption in SZ.

Adequate brain vascularization and BBB formation is crucial for proper brain development and functioning. Brain endothelium secretes and transports factors and nutrients that are important for neurogenesis, neuron activity and neuroprotection, among others. Therefore an intrinsic failure in these endothelial functions could be altering brain development at early stages, as well as interfering with brain recovery after insults [69–74].

Although the endothelial differentiation protocol used in this article has been lately questioned for the expression of epithelial genes [75], here we corroborated the expression of classical endothelial genes and BBB-genes, as was shown by the original authors [32, 76]. Moreover, we show that the expression of those factors is even higher in BEC than their expression in the commonly used BBB cell line (HCMEC/D3). In Ctrl BEC, VEGFA stimulations generates an angiogenic response and they also present high TEER values when cultured as monolayers. Therefore, we are confident on our results and propose that this differentiation protocol is useful for the evaluation of angiogenesis and BBB features in SZP hiPSC-derived BEC.

Our results set the ground for future studies on how the brain endothelium may modulate maturation and maintenance the of neuronal networks via the production and release of these soluble factors and point to the importance of the neurovascular niche in the CNS when analyzing the SZ phenotype. As increasing molecular data has been obtained from patients with SZ and new technologies allow us to navigate the otherwise unknown cerebral development. It is clear that SZ not only present specific neuronal alterations but may be the result of a greater combination of intrinsic deficiencies in the cellular function of glia and BEC. More research is needed to evaluate the potential implications of combining all this cell types in a neurovascular reconstruction of the SZ brain.

## Acknowledgements

We thank Dra. Carol San Martin for kindly donating HCMEC/D3 line. We are grateful to all the Steven Rehen’s laboratory crew, especially to Dr. Livia Goto, Ismael Gomez, and Scarlett Mercante for technical support. We thank Sofía Puvogel for critical reading of the manuscript and Dr. Emiliano Molina for technical assistance. Intramural grants provided from the D’Or Institute for Research and Education (IDOR). Funding from ANID Fondecyt # 1190083 (VP) & Conicyt # 21150781 (BSC).

## References

1. Skene NG, Bryois J, Bakken TE, Breen G, Crowley JJ, Gaspar HA, et al. Genetic identification of brain cell types underlying schizophrenia. Nat Genet. 2018;50:825–833.

2. Potkin SG, Kane JM, Correll CU, Lindenmayer J-P, Agid O, Marder SR, et al. The neurobiology of treatment-resistant schizophrenia: paths to antipsychotic resistance and a roadmap for future research. Npj Schizophr. 2020;6:1.

3. Avramopoulos D. Recent Advances in the Genetics of Schizophrenia. Mol Neuropsychiatry. 2018;4:35–51.

4. Ripke S, Neale BM, Corvin A, Walters JTR, Farh KH, Holmans PA, et al. Biological insights from 108 schizophrenia-associated genetic loci. Nature. 2014;511:421– 427.

5. Stilo S, Forti M, Murray R. Environmental risk factors for schizophrenia: Implications for prevention. Neuropsychiatry (London). 2011;1:457–466.

6. Brown AS. The environment and susceptibility to schizophrenia. Prog Neurobiol. 2011;93:23–58.

7. Birnbaum R, Weinberger DR. Genetic insights into the neurodevelopmental origins of schizophrenia. Nat Rev Neurosci. 2017;18:727–740.

8. Costain G, Bassett AS. Clinical applications of schizophrenia genetics: genetic diagnosis, risk, and counseling in the molecular era. Appl Clin Genet. 2012;5:1– 18.

9. Insel TR. Rethinking schizophrenia. Nature. 2010;468:187–193.

10. Selemon LD, Zecevic N. Schizophrenia: a tale of two critical periods for prefrontal cortical development. Transl Psychiatry. 2015;5:e623–e623.

11. Richetto J, Meyer U. Epigenetic Modifications in Schizophrenia and Related Disorders: Molecular Scars of Environmental Exposures and Source of Phenotypic Variability. Biol Psychiatry. 2021;89:215–226.

12. Najjar S, Pahlajani S, De Sanctis V, Stern JNH, Najjar A, Chong D. Neurovascular Unit Dysfunction and Blood–Brain Barrier Hyperpermeability Contribute to Schizophrenia Neurobiology: A Theoretical Integration of Clinical and Experimental Evidence. Front Psychiatry. 2017;8:83.

13. Katsel P, Roussos P, Pletnikov M, Haroutunian V. Microvascular anomaly conditions in psychiatric disease. Schizophrenia – angiogenesis connection. Neurosci Biobehav Rev. 2017;77:327–339.

14. Bleuler E. Dementia praecox or the group of schizophrenias. Oxford, England: International Universities Press; 1950.

15. Saili KS, Zurlinden TJ, Schwab AJ, Silvin A, Baker NC, Hunter 3rd ES, et al. Blood-brain barrier development: Systems modeling and predictive toxicology. Birth Defects Res. 2017;109:1680–1710.

16. Schmidt-Kastner R, van Os J, Esquivel G, Steinbusch HWM, Rutten BPF. An environmental analysis of genes associated with schizophrenia: hypoxia and vascular factors as interacting elements in the neurodevelopmental model. Mol Psychiatry. 2012;17:1194–1205.

17. Pong S, Karmacharya R, Sofman M, Bishop JR, Lizano P. The Role of Brain Microvascular Endothelial Cell and Blood-Brain Barrier Dysfunction in Schizophrenia. Complex Psychiatry. 2020;6:30–46.

18. Greene C, Hanley N, Campbell M. Blood-brain barrier associated tight junction disruption is a hallmark feature of major psychiatric disorders. Transl Psychiatry. 2020;10.

19. Marín-Padilla M. The human brain intracerebral microvascular system: development and structure. Front Neuroanat. 2012;6:1–14.

20. Ben-Zvi A, Liebner S. Developmental regulation of barrier- and non-barrier blood vessels in the CNS. J Intern Med. 2021;1:1–16.

21. Daneman R, Prat A. The blood-brain barrier. Cold Spring Harb Perspect Biol. 2015;7:a020412.

22. Iadecola C. The Neurovascular Unit Coming of Age: A Journey through Neurovascular Coupling in Health and Disease. Neuron. 2017;96:17–42.

23. Brennand K, Savas JN, Kim Y, Tran N, Simone A, Hashimoto-Torii K, et al. Phenotypic differences in hiPSC NPCs derived from patients with schizophrenia. Mol Psychiatry. 2015;20:361–368.

24. Hoffman GE, Schrode N, Flaherty E, Brennand KJ. New considerations for hiPSC-based models of neuropsychiatric disorders. Mol Psychiatry. 2019;24:49–66.

25. Ardhanareeswaran K, Mariani J, Coppola G, Abyzov A, Vaccarino FM. Human induced pluripotent stem cells for modelling neurodevelopmental disorders. Nat Rev Neurol. 2017;13:265–278.

26. Das D, Feuer K, Wahbeh M, Avramopoulos D. Modeling Psychiatric Disorder Biology with Stem Cells. Curr Psychiatry Rep. 2020;22:24.

27. Moslem M, Olive J, Falk A. Stem cell models of schizophrenia, what have we learned and what is the potential? Schizophr Res. 2019;210:3–12.

28. Balan S, Toyoshima M, Yoshikawa T. Contribution of induced pluripotent stem cell technologies to the understanding of cellular phenotypes in schizophrenia. Neurobiol Dis. 2019;131:104162.

29. Casas BS, Vitória G, do Costa MN, Madeiro da Costa R, Trindade P, Maciel R, et al. hiPSC-derived neural stem cells from patients with schizophrenia induce an impaired angiogenesis. Transl Psychiatry. 2018;8:48.

30. Brennand KJ, Simone A, Jou J, Gelboin-Burkhart C, Tran N, Sangar S, et al. Modelling schizophrenia using human induced pluripotent stem cells. Nature. 2011;473:221–225.

31. Sochacki J, Devalle S, Reis M, de Moraes Maciel R, da Silveira Paulsen B, Brentani H, et al. Generation of iPS cell lines from schizophrenia patients using a non-integrative method. Stem Cell Res. 2016;17:97–101.

32. Qian T, Maguire SE, Canfield SG, Bao X, Olson WR, Shusta E V, et al. Directed differentiation of human pluripotent stem cells to blood-brain barrier endothelial cells. Sci Adv. 2017;3:48–50.

33. Stebbins MJ, Wilson HK, Canfield SG, Qian T, Palecek SP, Shusta E V. Differentiation and characterization of human pluripotent stem cell-derived brain microvascular endothelial cells. Methods. 2016;101:93–102.

34. Prieto C, Casas B, Falcón P, Villanueva A, Lois P, Lattus J, et al. Downregulation of the Netrin-1 Receptor UNC5b Underlies Increased Placental Angiogenesis in Human Gestational Diabetes Mellitus. Int J Mol Sci. 2019;20:1408.

35. Carpentier G, Martinelli M, Courty J, Cascone I. Angiogenesis Analyzer for ImageJ. 4th ImageJ User Dev. Conf. Proc., Mondorf-Les-Bains, Luxembourg; 2012. p. 198–201.

36. Takahashi H, Hattori S, Iwamatsu A, Takizawa H, Shibuya M. A Novel Snake Venom Vascular Endothelial Growth Factor (VEGF) Predominantly Induces Vascular Permeability through Preferential Signaling via VEGF Receptor-1. J Biol Chem. 2004;279:46304–46314.

37. Brash JT, Ruhrberg C, Fantin A. Evaluating Vascular Hyperpermeability-inducing Agents in the Skin with the Miles Assay. J Vis Exp. 2018;2018.

38. Schmidt-kastner R, Os J Van, Steinbusch HWM. Gene regulation by hypoxia and the neurodevelopmental origin of schizophrenia. Schizophr Res. 2006;84:253– 271.

39. Fulzele S, Pillai A. Decreased VEGF mRNA expression in the dorsolateral prefrontal cortex of schizophrenia subjects. Schizophr Res. 2009;115:372–373.

40. Lee B-H, Hong J-P, Hwang J-A, Ham B-J, Na K-S, Kim W-J, et al. Alterations in plasma vascular endothelial growth factor levels in patients with schizophrenia before and after treatment. Psychiatry Res. 2015;228:95–99.

41. Xiao W, Zhan Q, Ye F, Tang X, Li J, Dong H, et al. Baseline serum vascular endothelial growth factor levels predict treatment response to antipsychotic medication in patients with schizophrenia. Eur Neuropsychopharmacol. 2018;28:603–609.

42. Ye F, Zhan Q, Xiao W, Tang X, Li J, Dong H, et al. Altered serum levels of vascular endothelial growth factor in first-episode drug-naïve and chronic medicated schizophrenia. Psychiatry Res. 2018;264:361–365.

43. Mehlen P, Delloye-Bourgeois C, Chédotal A. Novel roles for Slits and netrins: axon guidance cues as anticancer targets? Nat Rev Cancer. 2011;11:188–197.

44. Podjaski C, Alvarez JI, Bourbonniere L, Larouche S, Terouz S, Bin JM, et al. Netrin 1 regulates blood–brain barrier function and neuroinflammation. Brain. 2015;138:1598–1612.

45. Prieto CP, Ortiz MC, Villanueva A, Villarroel C, Edwards SS, Elliott M, et al. Netrin-1 acts as a non-canonical angiogenic factor produced by human Wharton’s jelly mesenchymal stem cells (WJ-MSC). Stem Cell Res Ther. 2017;8:43.

46. Vafadari B, Salamian A, Kaczmarek L. MMP-9 in translation: from molecule to brain physiology, pathology, and therapy. J Neurochem. 2016;139:91–114.

47. Brilha S, Ong CWM, Weksler B, Romero N, Couraud P-O, Friedland JS. Matrix metalloproteinase-9 activity and a downregulated Hedgehog pathway impair blood-brain barrier function in an *in vitro* model of CNS tuberculosis. Sci Rep. 2017;7:16031.

48. Kadry H, Noorani B, Cucullo L. A blood–brain barrier overview on structure, function, impairment, and biomarkers of integrity. Fluids Barriers CNS. 2020;17:69.

49. Huang G, Osorio D, Guan J, Ji G, Cai JJ. Overdispersed gene expression in schizophrenia. Npj Schizophr. 2020;6:9.

50. Krock BL, Skuli N, Simon MC. Hypoxia-induced angiogenesis: good and evil. Genes Cancer. 2011;2:1117–1133.

51. Schmidt-Kastner R, Guloksuz S, Kietzmann T, van Os J, Rutten BPF. Analysis of GWAS-Derived Schizophrenia Genes for Links to Ischemia-Hypoxia Response of the Brain. Front Psychiatry. 2020;11:1–9.

52. Tornavaca O, Chia M, Dufton N, Almagro LO, Conway DE, Randi AM, et al. ZO-1 controls endothelial adherens junctions, cell–cell tension, angiogenesis, and barrier formation. J Cell Biol. 2015;208:821–838.

53. Dev KK, Henley JM. The schizophrenic faces of PICK1. Trends Pharmacol Sci. 2006;27:574–579.

54. Li C, Tao R, Qin W, Zheng Y, He G, Shi Y, et al. Positive association between PDLIM5 and schizophrenia in the Chinese Han population. Int J Neuropsychopharmacol. 2008;11:27–34.

55. Leduc-Galindo D, Qvist P, Tóth AE, Fryland T, Nielsen MS, Børglum AD, et al. The effect of hypoxia on ZEB1 expression in a mimetic system of the blood-brain barrier. Microvasc Res. 2019;122:131–135.

56. Li Y, Xia Y, Zhu H, Luu E, Huang G, Sun Y, et al. Investigation of Neurodevelopmental Deficits of 22 q11.2 Deletion Syndrome with a Patient-iPSC-Derived Blood–Brain Barrier Model. Cells. 2021;10:2576.

57. Pong S, Lizano P, Karmacharya R. Investigating Blood-Brain Barrier Dysfunction in Schizophrenia Using Brain Microvascular Endothelial Cells Derived From Patient-Specific Stem Cells. Biol Psychiatry. 2020;87:S189–S190.

58. Crockett AM, Ryan SK, Vásquez AH, Canning C, Kanyuch N, Kebir H, et al. Disruption of the blood–brain barrier in 22q11.2 deletion syndrome. Brain. 2021;144:1351–1360.

59. Hashimoto K, Shimizu E, Komatsu N, Nakazato M, Okamura N, Watanabe H, et al. Increased levels of serum basic fibroblast growth factor in schizophrenia. Psychiatry Res. 2003;120:211–218.

60. Greene C, Kealy J, Humphries MM, Gong Y, Hou J, Hudson N, et al. Dose-dependent expression of claudin-5 is a modifying factor in schizophrenia. Mol Psychiatry. 2018;23:2156–2166.

61. Kis B, Chen L, Ueta Y, Busija DW. Autocrine peptide mediators of cerebral endothelial cells and their role in the regulation of blood–brain barrier. Peptides. 2006;27:211–222.

62. Vermeer PD, Denker J, Estin M, Moninger TO, Keshavjee S, Karp P, et al. MMP9 modulates tight junction integrity and cell viability in human airway epithelia. Am J Physiol Cell Mol Physiol. 2009;296:L751–L762.

63. Domenici E, Willé DR, Tozzi F, Prokopenko I, Miller S, McKeown A, et al. Plasma protein biomarkers for depression and schizophrenia by multi analyte profiling of case-control collections. PLoS One. 2010;5:e9166.

64. Yamamori H, Hashimoto R, Ishima T, Kishi F, Yasuda Y, Ohi K, et al. Plasma levels of mature brain-derived neurotrophic factor (BDNF) and matrix metalloproteinase-9 (MMP-9) in treatment-resistant schizophrenia treated with clozapine. Neurosci Lett. 2013;556:37–41.

65. Chang S-H, Chiang S-Y, Chiu C-C, Tsai C-C, Tsai H-H, Huang C-Y, et al. Expression of anti-cardiolipin antibodies and inflammatory associated factors in patients with schizophrenia. Psychiatry Res. 2011;187:341–346.

66. Han H, He X, Tang J, Liu W, Liu K, Zhang J, et al. The C(−1562)T polymorphism of matrix metalloproteinase-9 gene is associated with schizophrenia in China. Psychiatry Res. 2011;190:163–164.

67. Rybakowski JK, Skibinska M, Kapelski P, Kaczmarek L, Hauser J. Functional polymorphism of the matrix metalloproteinase-9 (MMP-9) gene in schizophrenia. Schizophr Res. 2009;109:90–93.

68. Wiera G, Wozniak G, Bajor M, Kaczmarek L, Mozrzymas JW. Maintenance of long-term potentiation in hippocampal mossy fiber-CA3 pathway requires fine-tuned MMP-9 proteolytic activity. Hippocampus. 2013;23:529–543.

69. Glasgow SD, Ruthazer ES, Kennedy TE. Guiding synaptic plasticity: Novel roles for netrin-1 in synaptic plasticity and memory formation in the adult brain. J Physiol. 2021;599:493–505.

70. Glasgow SD, Labrecque S, Beamish I V., Aufmkolk S, Gibon J, Han D, et al. Activity-Dependent Netrin-1 Secretion Drives Synaptic Insertion of GluA1-Containing AMPA Receptors in the Hippocampus. Cell Rep. 2018;25:168-182.e6.

71. Bayas A, Hummel V, Kallmann BA, Karch C, Toyka K V., Rieckmann P. Human cerebral endothelial cells are a potential source for bioactive BDNF. Cytokine. 2002;19:55–58.

72. Nakahashi T, Fujimura H, Altar CA, Li J, Kambayashi JI, Tandon NN, et al. Vascular endothelial cells synthesize and secrete brain-derived neurotrophic factor. FEBS Lett. 2000;470:113–117.

73. Yi H, Hu J, Qian J, Hackam AS. Expression of brain-derived neurotrophic factor is regulated by the Wnt signaling pathway. Neuroreport. 2012;23:189–194.

74. Wu H, Lu D, Jiang H, Xiong Y, Qu C, Li B, et al. Simvastatin-mediated upregulation of VEGF and BDNF, activation of the PI3K/Akt pathway, and increase of neurogenesis are associated with therapeutic improvement after traumatic brain injury. J Neurotrauma. 2008;25:130–139.

75. Lu TM, Houghton S, Magdeldin T, Durán JGB, Minotti AP, Snead A, et al. Pluripotent stem cell-derived epithelium misidentified as brain microvascular endothelium requires ETS factors to acquire vascular fate. Proc Natl Acad Sci. 2021;118:e2016950118.

76. Lippmann ES, Azarin SM, Palecek SP, Shusta E V. Commentary on human pluripotent stem cell-based blood–brain barrier models. Fluids Barriers CNS. 2020;17:64.

