## Supplementary Material for "Schizophrenia-derived hiPSC brain microvascular endothelial cells show impairments in angiogenesis and blood-brain barrier function"

**Supplementary Information**


Supplementary Table 1
Supplementary Table 2
Supplementary Figure 1
Supplementary Figure 2
Supplementary Figure 3
Supplementary Figure 4

| Identification | Cell line | Group | Gender | Age | Cell Source | Reprogramming Technique |
|---|---|---|---|---|---|---|
| Ctrl #1 | GM23279A | Control | M | 37 | Fibroblast | Lentiviral vector (OCT4, SOX2, KLF4, MYC, LIN28) |
| Ctrl #2 | CF1 | Control | M | 31 | Fibroblast | Cytotune 1.0 kit (ThermoFisher) |
| Ctrl #3 | CF2 | Control | F | 36 | Fibroblast | Cytotune 1.0 kit (ThermoFisher) |
| SZP #1 | GM23760B | Schizophrenia | M | 26 | Fibroblast | Lentiviral vector (OCT4, SOX2, KLF4, MYC, LIN28) |
| SZP #2 | GM23761B | Schizophrenia | F | 27 | Fibroblast | Lentiviral vector (OCT4, SOX2, KLF4, MYC, LIN28) |
| SZP #3 | GM23762B | Schizophrenia | M | 23 | Fibroblast | Lentiviral vector (OCT4, SOX2, KLF4, MYC, LIN28) |
| SZP #4 | EZQ4 | Schizophrenia | M | 42 | Fibroblast | Cytotune 2.0 kit (ThermoFisher) |
| SZP #5 | EZQ9 | Schizophrenia | M | 44 | Fibroblast | Cytotune 2.0 kit (ThermoFisher) |

**Supplementary Table 1.** Sample information. Each cell line used was obtained from fibroblast donated from healthy subjects (Control group) or patients diagnosed with schizophrenia (Schizophrenia group).

| Gene | Fw sequence | Rv sequence |
|---|---|---|
| **18S** | GGGCCCGAAGCGTTTACTTT | TTGCGCCGGTCCAAGAATTT |
| **PCAM-1 (CDH5)** | TTTTCACAGGGCGGGTTTCT | TGACCCTCAAGGCTTGAACA |
| **CLDN5** | GCGGGTGTCAGACTGAGGATT | GCCCTGCCGATGGAGTAAAGA |
| **CDH5** | AGAATGACAATGCCCCGGAGTT | GATGTTGGCCGTGTTATCGTGA |
| **FLT1** | ATCACTCAGCGCATGGCAAT | TCTCCTTCCGTCGGCATTTT |
| **GAPDH** | CAAGAAGGTGGTGAAGCAGGC | CCACCACCCTGTTGCTGTAG |
| **GLUT-1** | TGCCTGAAGTCGCACAGTGAA | AGGGCAGCTTGACAGCTCATT |
| **HIF1a** | AGCCGCTGGAGACACAATCAT | TCGAAGTGGCTTTGGCGTTT |
| **KDR** | TCATGCACGGCATCTGGGAAT | GCACAGCCAAGAACACTGCAT |
| **NRP1** | AGCCTGCAACTTGGGAAACT | TGGTTACCAGGCGGATGTTT |
| **NRP2** | TCTGGCCGGATTGCTAATGA | TGGCCGTGAGCATGGTTAAA |
| **OCLN** | TGCTTCAGCATCAGGATGGAAGTG | GCAGAATCATGAGCTGCCAGGAAA |
| **P-gp (MDR-1)** | AACAGTCCAGCTGATGCAGA | TTCACGGCCATAGCGAATGT |
| **uPA** | GGAAAACCTCATCCTACACAA | CGGATCTTCAGCAAGGCAATG |
| **vWF** | TGCCAGAGCCTGCACATCAAT | CCACTGGCTGTTTCGGCAAAT |
| **ZO-1** | TCACGCAGTTACGAGCAAGT | GAGGCAGTGGTTTGGTGTTT |
| **NTN1** | TGCAAGAAGGACTATGCCGTC | GCTCGTGCCCTGCTTATACAC |
| **NEO1** | GCTTCATCAAATTGACGTGGCGGA | AGATGTACACGGTCGCTGGCATTA |
| **DCC** | GGGGCCACTCTCTGATCCTA | TGCATTTGTCCAATTGGCGG |
| **UNC5b** | GGGCTGGAGGATTACTGGTG | TGCAGGAGAACCTCATGGTC |
| **ADORA2B** | GTCTGCCTTGTTTATGGTGGAA | GAGGTCACCTTCCTGGCAAC |

**Supplementary Table 2.** Primers used for qRT-PCR.

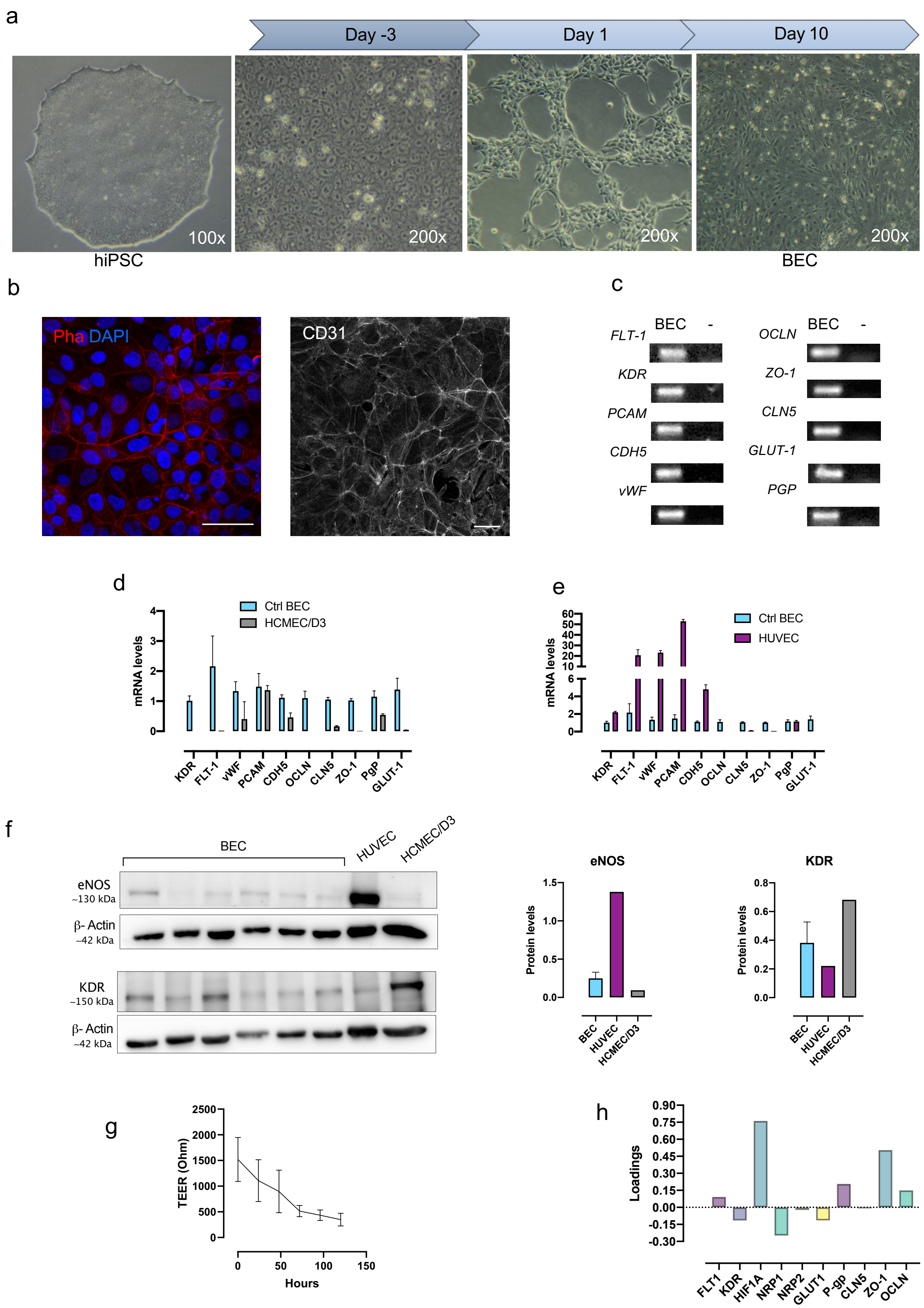


**Supplementary Figure 1. hiPSC derivation to BEC.** hiPSC were differentiated to microvascular brain endothelial like cells following Qian et al protocol. **a** Representative image of cells at mentioned days of the differentiation protocol. **b** Representative immunostaining of BEC showing clear cobblestone morphology (left) and CD31 expression in plasma membrane (right). Bar= 50 µm. Phalloidin staining was used to asses actin (Pha, red), DAPI was used for nuclei staining (blue). **c** Expression of classic endothelial and BBB-specific genes in BEC. **d-e** mRNA levels of classic endothelial and BBB-specific genes was assessed by qPCR in Ctrl BEC, HCMEC/D3 (d) and HUVEC (e); *B2M* was used as housekeeping gene. Data is expressed as mean ± SD. **f** Representative membrane of WB of eNOS and KDR (left) and band quantification by densitometry (right) performed on BEC, HUVEC and HCMEC/D3. **g** TEER measurement in Ctrl BEC up to 6 days after differentiation (N=3).**h** Gene loadings for PC 1 (Figure 1d).

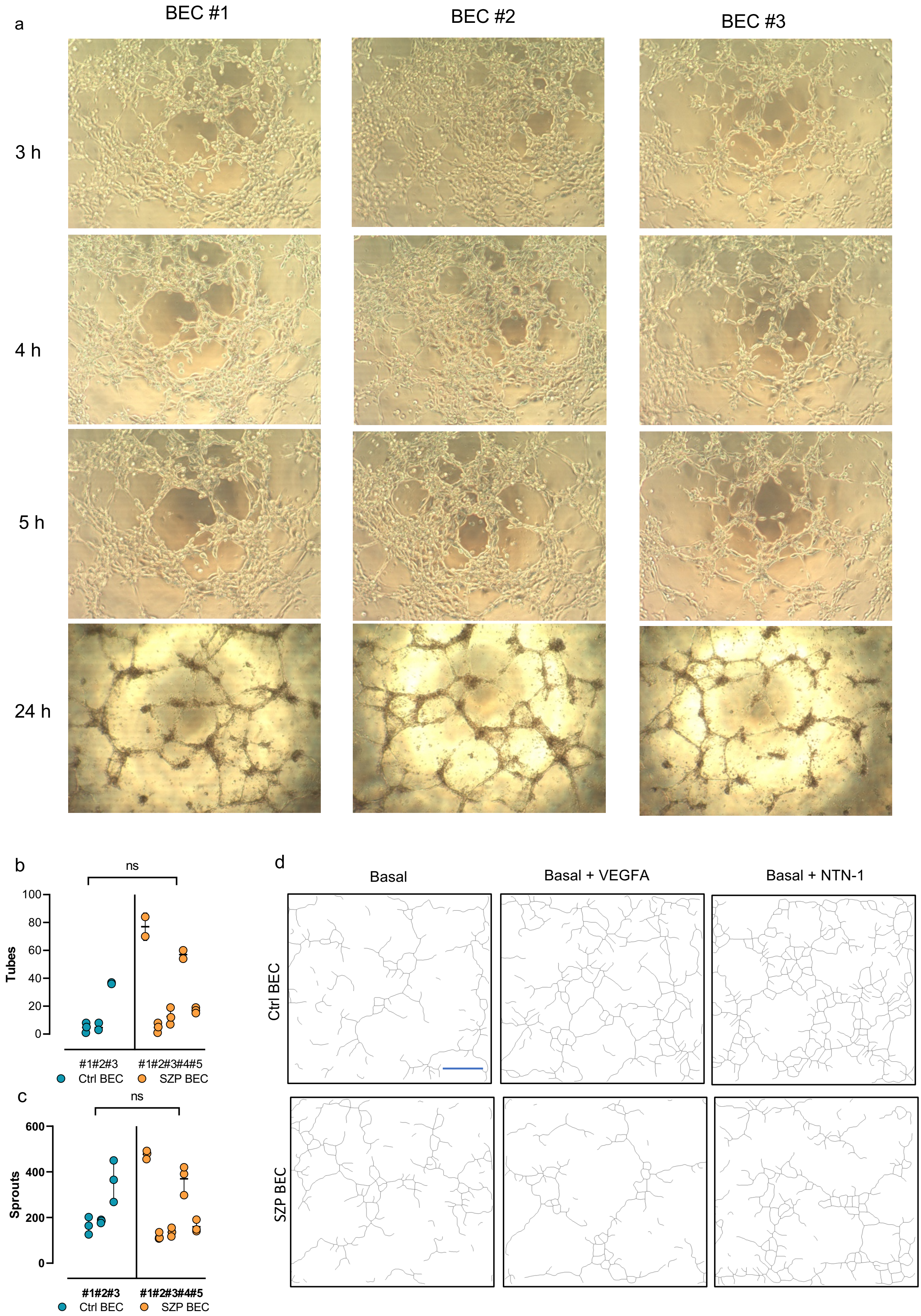
a
BEC #1
BEC #2
BEC #3
3 h
4 h
5 h
24 h
b
ns
Tubes
100
80
60
40
20
0
#1#2#3
#1#2#3#4#5
Ctrl BEC
SZP BEC
c
ns
Sprouts
600
400
200
0
#1#2#3
#1#2#3#4#5
Ctrl BEC
SZP BEC
d
Basal
Basal + VEGFA
Basal + NTN-1
Ctrl BEC
SZP BEC

**Supplementary Figure 2**. **Tube formation assay on BEC. a** Representative images of tube formation assay performed to three Ctrl BEC. Pothographs were taken after 3, 4, 5 and 24 h of seeding. Tubes start forming after 3 h and reach a maximum after 4 h. At 5 h some structures are lost. Images of 24 h are at half magnification and show great loss of tubular structures. **b-c** Quantification of tube formation assay of Ctrl and SZP BEC, when incubating BEC with Basal media (hESFM + B27). Graphs show number of sprouts **(b)** and tubes **(c)** for each cell donor. There was no significant difference (ns) between groups in any case, according to Nested t-test. **d** Descriptive segmentation of images on Figure 2a, performed by Angiogenesis Analyzer software in order to quantify tubes and sprouts. Scale bar = 30 µm.

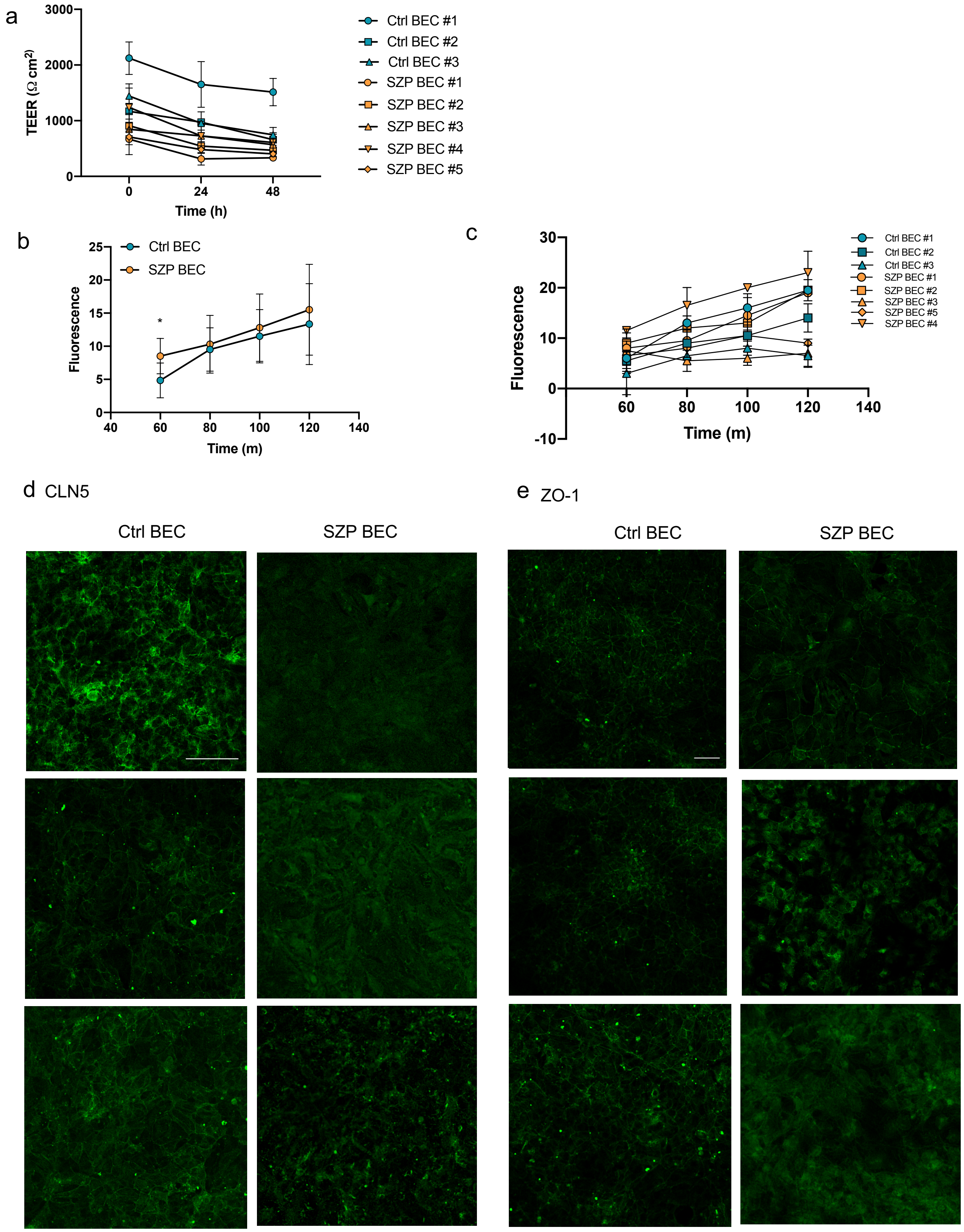


**Supplementary Figure 3. a** TEER was measured the first day of BEC differentiation (time=0 h) and after 24 and 48 h of culture. Data is shown as mean ± SD. Each cell donor is identified with correlative numbers in the graph (Ctrl BEC: #1, #2, #3; SZP BEC: #1, #2, #3, #4, #5. **b** Dextran (fluorescence) permeability was assessed in Ctrl and SZP BEC within a two hour period. Data is shown as mean ± SD and includes all biological replicates of Ctrl and SZP group. * $p < 0.05$ according to two-way ANOVA. **c** Graph shows dextran (fluorescence) permeability for each cell donor of (b) and is identified with correlative numbers in the graph (Ctrl BEC: #1, #2, #3; SZP BEC: #1, #2, #3, #4, #5). **d-e** Representative immunofluorescences of CLN5 (d) and ZO-1 (e) in Ctrl and SZP BEC; bar= 50µm.

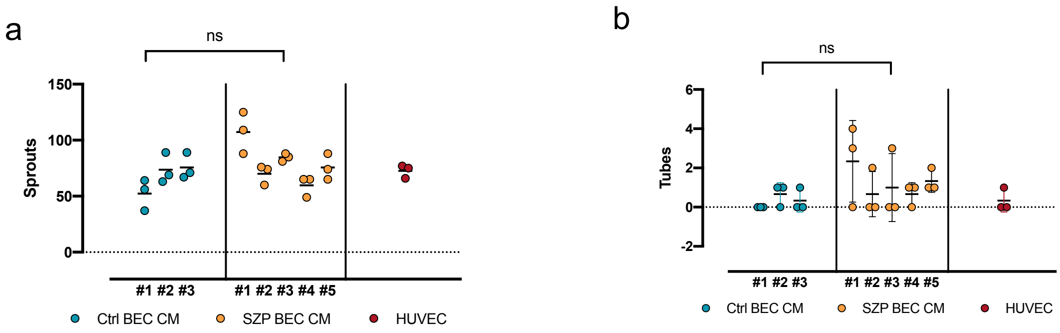


**Supplementary Figure 4.** Quantification of average number of sprouts (a) and tubes (b) generated after stimulation of HCMEC/D3 Ctrl BEC CM, SZP BEC CM and HUVEC CM. Stimulation was performed in triplicate for each cell donor CM, identified with correlative numbers in the graph (Ctrl BEC CM #1-3 and SZP BEC CM #1-5).

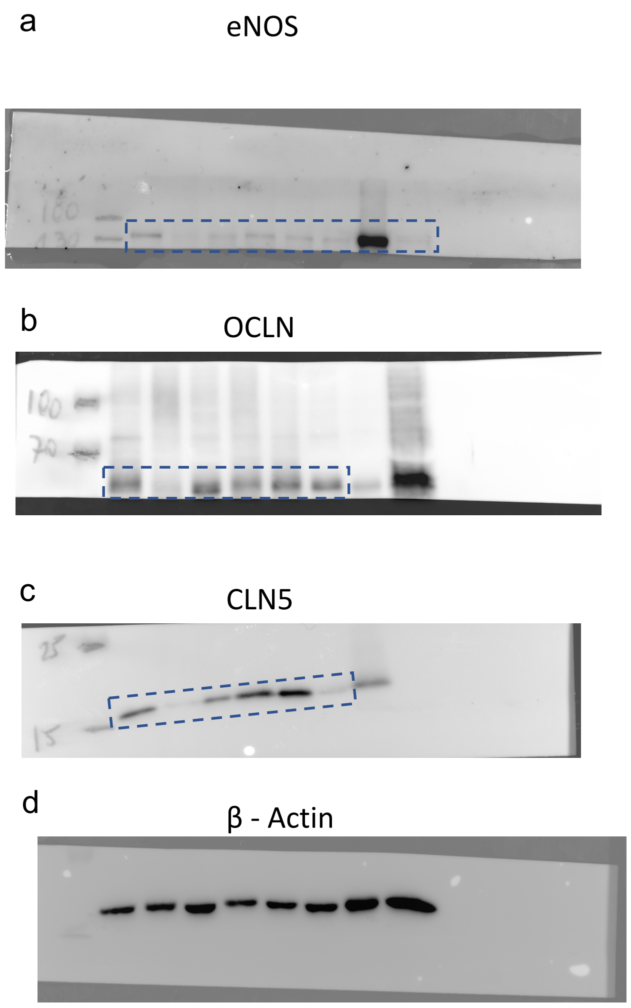


**Supplementary Figure 5.** Complete images of westernblot membranes. a-d were transferred from the same gel. e-f were transferred from the same gel. Lanes 1-3: Ctrl BEC; Lanes 4-6: SZP BEC; Lane 7: HUVEC; Lane 8: HCMEC/D3. Blue dashed boxes show bands that were shown in Supplementary Figure 1 and Figure 3.
